# Transcriptomic and histological characterization of telocytes in the human dorsal root ganglion

**DOI:** 10.1101/2024.09.24.614693

**Authors:** Rainer V Haberberger, Dusan Matusica, Stephanie Shiers, Ishwarya Sankaranarayanan, Theodore J Price

## Abstract

Telocytes are interstitial cells with long processes that cover distances in tissues and likely coordinate interacts with other cell types. Though present in central and peripheral neuronal tissues, their role remains unclear. Dorsal root ganglia (DRG) house pseudounipolar afferent neurons responsible for signals such as temperature, proprioception and nociception. This study aimed to investigate the presence and function of telocytes in human DRG by investigating their transcriptional profile, location and ultrastructure.

Sequencing data revealed CD34 and PDGFRA expressing cells comprise roughly 1.5-3% of DRG cells. Combined expression of *CD34* and *PDGFRA* is a putative marker gene set for telocytes. Further analysis identified nine subclusters with enriched cluster-specific genes. KEGG and GO pathway analysis suggested vascular, immune and connective tissue associated putative telocyte subtypes. Over 3000 potential receptor-ligand interactions between sensory neurons and these *CD34* and *PDGFRA* expressing putative telocytes were identified using a ligand-receptors interactome platform. Immunohisto-chemistry showed CD34+ telocytes in the endoneural space of DRGs, next to neuron-satellite complexes, in perivascular spaces and in the endoneural space between nerve fibre bundles, consistent with pathway analysis. Transmission electron microscopy (TEM) confirmed their location identifying characteristic elongated nucleus, long and thin telopods containing vesicles, surrounded by a basal lamina. This is the first study that provides gene expression analysis of telocytes in complex human tissue such as the DRG, highlighting functional differences based on tissue location with no significant ultrastructural variation.

**Key points:**

- uman DRGs contain *CD34/PDGFRA*+ putative telocytes that can be subgrouped in nine clusters based on gene expression
- The DRGs contain CD34+ telocytes in close proximity to neuron-satellite complexes, blood vessels and nerve fibre bundles.
- The basic features of telocytes with elongated nuclei and thin telopods do not differ between locations.

## 1 Introduction

Telocytes are interstitial, stromal cells found in connective tissues across many organs of the human body (see reviews (S. M. Cretoiu & Popescu, 2014; Kondo & Kaestner, 2019)). They possess a characteristic structure with long cellular processes (telopods) that can show dilations (podoms) with segments between podoms termed podomers (Xiao & Bei, 2016). Telocytes build loose networks with connecting telopods that can extend over 100 μm in length (S. M. Cretoiu & Popescu, 2014) allowing connection with various other cell types (S. M. Cretoiu & Popescu, 2014; Kondo & Kaestner, 2019). They can be labelled immunohistochemically with markers such as the hematopoietic stem cell marker CD34 or the platelet-derived growth factor (PDGF) receptor alpha (Xiao & Bei, 2016) although the immunohistochemical profile may vary across tissues (Diaz-Flores et al., 2020).

Telocytes have been described in many human organs (Kondo & Kaestner, 2019) such as the gastrointestinal tract (GIT) (Milia et al., 2013; Pieri, Vannucchi, & Faussone-Pellegrini, 2008), heart (D. Cretoiu, Hummel, Zimmermann, Gherghiceanu, & Popescu, 2014; Gherghiceanu & Popescu, 2012), lung (Manetti et al., 2014), urinary system (Vannucchi, Traini, Guasti, Del Popolo, & Faussone-Pellegrini, 2014), skin (Manole, Gherghiceanu, Ceafalan, & Hinescu, 2022) and the peripheral nervous system including enteric and parasympathetic ganglia (Diaz-Flores et al., 2020; Pieri et al., 2008; Rusu et al., 2016; Vandecasteele et al., 2017) and have been investigated in those organs under normal and pathological conditions including Crohn’s disease (Milia et al., 2013), atherosclerosis (Xu, Tian, Qiao, & Zheng, 2021) and urinary system pathologies (Sanches et al., 2021; Traini et al., 2018; Wishahi, Hafiz, Wishahy, & Badawy, 2021). However, the presence and ultrastructure of telocytes in human dorsal root ganglia (DRG) remain unknown. Recent transcriptomic studies have identified gene expression markers for telocytes which include *CD34*, *PDGFRA* and, to with less specificity, *DCN* (Guo et al., 2024) but gene expression in these cells has not been investigated at all in any human nervous system tissue.

Dorsal root ganglia (DRG) house the cell bodies of pseudounipolar sensory neurons responsible for transmitting nociception, temperature, itch or mechanical stimulation signals (Haberberger, Barry, Dominguez, & Matusica, 2019). These sensory neurons form a complex with satellite glia cells. Adjacent capillaries supply blood to meet energy demands, while Schwan cells envelop neuronal processes. Mammalian DRGs harbor several other cell types in addition to the neuron-satellite cell complex including macrophages, fibroblasts, and pericytes (Haberberger et al., 2019; Haberberger, Kuramatilake, Barry, & Matusica, 2023; Tavares-Ferreira et al., 2022). Recent studies explored the cell types in human DRGs via RNA-sequencing (RNA-seq) and subdivided the cells based on transcript enrichment into satellite glia cells (SGC) (Avraham et al., 2022), sensory neurons, and other non-neuronal cells (Bhuiyan et al., 2024; Jung et al., 2023; Nguyen, von Buchholtz, Reker, Ryba, & Davidson, 2021; Tavares-Ferreira et al., 2022). Despite this, few studies have focussed on analysis of non-neuronal cell types in human DRG (Avraham et al., 2022; Scaravilli, Giometto, Chimelli, & Sinclair, 1991). Using a whole cell RNA-seq technique we identified a subset of non-neuronal, non-glial cells that possess a transcriptional identity consistent with telocytes. We confirmed the presences of these cells in the human DRG based on immunohistochemical characteristics and ultrastructural morphology. We propose that telocytes in the human DRG may play a key role in communication between the vasculature and neuron-glial units. In this study we demonstrate the presence, transcriptional profile, and ultrastructure of telocytes in the human DRG, and propose functional roles for how these cells may interact with local sensory neurons, glial cells and blood vessels within these ganglia.

## 2 Material & Methods

### 2.1 Human tissue

All human tissue procurement procedures were approved by the Institutional Review Boards at the University of Texas at Dallas. Human lumbar DRGs were procured from three organ donors through a collaboration with the Southwest Transplant Alliance. DRGs were recovered using a ventral approach as previously described (Valtcheva et al., 2016). Donor medical history was provided by the Southwest Transplant Alliance and includes medical details from the donor’s family and hospital records. Donor demographics, medical history, and DRG level details are provided in **Supplementary Table 1.** For DRGs that were used for histological studies, upon removal from the body, the DRGs were cut in half, placed in 4% paraformaldehyde (PFA) for 48 hours at 4°C, and then transferred to Phosphate Buffered Saline (PBS) and shipped on ice to the University of Adelaide in Australia for histological analyses. For DRGs used for RNA-seq methods are described below.

### 2.2 Single-Cell RNA Sequencing

Single-cell RNA sequencing from human DRGs was performed as previously described (Hou et al., 2024). Human DRGs were surgically extracted from donors and placed in chilled, aerated artificial cerebrospinal fluid (aCSF). The aCSF solution contained 93 mM N-Methyl-D-glucamine (NMDG; Sigma-Aldrich), HCl (12 N; Fisher), KCl (Sigma-Aldrich), NaH2PO4 (Sigma-Aldrich), NaHCO_3_ (Sigma-Aldrich), HEPES (Sigma-Aldrich), D-(+)-Glucose (Sigma-Aldrich), L-Ascorbic acid (Sigma-Aldrich), Thiourea (Sigma-Aldrich), Na+ pyruvate (Sigma-Aldrich), MgSO_4_ (2 M; Fisher), CaCl_2_ dihydrate (Sigma-Aldrich), and N-acetylcysteine (Sigma-Aldrich). The DRGs were transferred to a sterile petri dish on ice and trimmed to remove connective tissue and fat, using forceps and Bonn scissors. The dural coats (perineurium and epineurium) were carefully removed, isolating the ganglia bodies. These were further divided into approximately 1 mm thick sections using Bonn scissors. The tissue fragments were placed in 5 mL of a prewarmed enzyme mix containing Stemxyme 1, Collagenase/Neutral Protease Dispase, and Deoxyribonuclease I (DNase I) in sterile filtered Hank’s Balanced Salt Solution (HBSS). The DRG-enzyme mixture was placed in a shaking water bath. It was gently triturated every 25 minutes using a sterile fire-polished glass Pasteur pipette until the solution turned cloudy and the tissue chunks passed smoothly through the pipette without resistance. Following enzymatic digestion, the dissociated DRGs were passed through a 100 µm cell strainer to remove debris and achieve a uniform cell suspension. The DRG cells were further isolated by layering the cell suspension on a 10% Bovine Serum Albumin (BSA) solution prepared in sterile HBSS. The BSA gradient was then centrifuged at 300 x g for 10 minutes, resulting in the isolation of DRG cells. The supernatant was discarded, and the cell pellet was immediately fixed using the 10X Chromium Fixed RNA Profiling kit. The cells were fixed for 17 hours at 4°C, followed by incubation with the 10X Fixed RNA Feature Barcode kit for 16 hours. The remainder of the library preparation was conducted according to the manufacturer’s protocol. The samples were sequenced using a NextSeq2000 at the genome core facility at the University of Texas at Dallas. Sequencing data were processed and mapped to the human genome (GRCh38) using 10X Genomics Cell Ranger v7.

Data analysis for single-cell RNA sequencing was conducted using the Seurat integration workflow (Stuart et al., 2019). To ensure data quality, only cells with less than 5% mitochondrial gene expression were included. The analysis workflow involved normalizing the data and selecting the top 2000 most variable features. Following data scaling, standard clustering techniques were applied, and the results were visualized using Uniform Manifold Approximation and Projection (UMAP) with a resolution parameter set to 1.

Next, we subset Telocytes from our single-cell database that met the conditions *CD34* > 1 and *PDGFRA* > 1. Cluster-specific markers were identified using the Wilcoxon rank-sum test implemented in Seurat. We ranked the marker gene list of telocytes based on Log2-fold change. Using the Sensoryomic web tool (https://sensoryomics.shinyapps.io/Interactome/), we performed an interactome analysis (Wangzhou et al., 2021) of Telocyte marker genes and the human DRG neuron expression data (Bhuiyan et al., 2024; Tavares-Ferreira et al., 2022). The analysis focused on telocyte-receptor interactions with hDRG ligands and vice versa, identifying the top 50 receptor-ligand interactions.

The top 150 genes of each of the reclustered telocyte subclusters were analysed by association with the Gene Ontology (GO) database and the Kyoto Encyclopedia of Genes and Genomes (KEGG) using the online database STRING (http://STRING-db.org). The database was also used for protein-protein network analysis.

### 2.3 Multiple labelling immunohistochemistry (IHC)

Upon receipt from UTDallas, one half of an individual DRG was embedded in paraffin (Paraplast, Leica) using an automatic paraffin embedding system (HistoPearl, Leica). Sections (4 μM thickness) were cut, mounted on glass slides for haematoxylin and eosin (H&E) staining and on polyethylenimine (PEI)-coated slides for immunohistochemistry, dried overnight, deparaffinized using xylene and decreasing concentrations of ethanol and rehydrated. Sections were stained using routine H&E staining to evaluate the quality of the tissue and subsequently a second set of sections processed for multiple labelling immunohistochemistry.

For multiple labelling immunohistochemistry 10 mM citrate buffer pH 6 or Tris/EDTA pH 9 were used for antigen retrieval, followed by washing in PBS and multiple labelling IHC. Before incubation with the primary antiserum or combination of antisera for 48h at room temperature, unspecific binding sites were blocked using normal donkey serum. Incubation with the primary antisera was followed by washing in PBS and 2h incubation with secondary antisera in combination with the nuclear stain 4′,6-diamidino-2-phenylindole (DAPI, Table 1). After an additional washing step in PBS, sections were covered with buffered glycerol pH 8.6, coverslipped and sealed with nail varnish. A minimum of three technical replicates from each of the DRGs were investigated for each combination of antisera. The monoclonal CD34 antibody is directed against a human extracellular class II epitope. Binding specificity was checked by positive staining of Kaposi sarcoma. The S-100 polyclonal antiserum is directed against purified S-100 protein from bovine brain and shows S-100 protein in human nervous system (Migheli et al., 1999). The monoclonal Nf200 antibody is directed against intermediate filament, pig neurofilament 200 kDa H-subunit. It has been used in human peripheral neuronal tissue (Chang et al., 2018; Henry, Luo, & Levinson, 2012). In humans, NF200 is a marker for all DRG neurons (Haberberger et al., 2019; Rostock, Schrenk-Siemens, Pohle, & Siemens, 2018). High resolution images for entire sections of human DRG were acquired using a Slide Scanner (20x objective, Olympus VS200) and sections further investigated using confocal microscopy (Zeiss LSM 880 Fast Airyscan, Leica Mica).

**Table 1.** Primary, secondary antibodies and DAPI.

| Antigen | Host | Dilution | Supplier |
| --- | --- | --- | --- |
| CD34 | Mouse, clone QBEN10 | 1:600 | Thermofisher |
| S100 | Rabbit | 1:2000 | Sigma |
| NF200 | Mouse, clone N52 | 1:2000 | Sigma |
| DAPI (4',6-diamidino-2-phenylindole) |  | 1:1000 | Invitrogen |
| Antigen | Host | Dilution | Supplier |
| Mouse IgG-CY3 | Donkey | 1:100 | Jackson |
| Rabbit IgG-CY3 | Donkey | 1:100 | Jackson |
| Rabbit IgG-CY5 | Donkey | 1:100 | Jackson |

### 2.4 Electron microscopy

The second half of the DRG was postfixed in 10% formalin for 48h, washed in PBS and stored at 4°C until further processing. The tissues were subsequently cut into smaller pieces of about 1 mm^3^ in size, fixed in 2.5% glutaraldehyde 2.5 % in phosphate buffer, pH 7.4 at 4° C for 24h, washed in PBS and transferred into 2% aqueous osmium tetroxide solution for 1h. Then the samples were dehydrated in graded series of ethanol and embedded in TAAB epon araldite embedding medium at 60° C for 48h. Semi-thin sections with a thickness of 500 nm were prepared and stained with 0.05% toluidine blue in borate buffer for 1 min. Ultrathin sections of 70-90 nm thickness were cut using a ultramicrotome (Leica), stained with 4% uranyl acetate and Reynolds lead citrate for 8 min and examined using an electron microscope (FEI Tecnai 120kV Spirit). Images were captured using an AMT Camera with AMT_V7.0.1 software. Three to six sections from three different areas were investigated.

## 3. Results

### 3.1 *In-silico* identification of *CD34* and *PDGFRA* expressing cells in human DRG

Since it is known that telocytes express both *CD34* and *PDGFRA* we first sought to identify cells expressing both of these markers in a previously published harmonized atlas of DRG cell types. In this dataset of 34,693 non-neuronal single nuclei, we observed 767 cells that co-expressed *CD34* and *PDGFRA* (Bhuiyan et al., 2024). To further confirm this finding, we used a whole cell, single cell RNA-sequencing dataset we have previously described that has the advantage of being deeply sequenced and contains cytoplasmic mRNAs, giving a more thorough picture of gene expression for any given cell. Out of 41,678 cells sequenced, we identified 1,287 cells that expressed both *CD34* and *PDGFRA*, making these putative telocytes (**Fig 1A and B**). We then generated a read count matrix for all 1,287 cells creating a list of genes expressed in these cells on a per cell basis (**Supplementary Table 2**). From this we re-clustered the cells and identified 9 subclusters that express both *CD34* and *PDGFRA* and we identified gene expression markers that were enriched in each cell population (**Supplementary Table 3**). These 9 subclusters of cells also expressed the telocyte marker *DCN* (Guo et al., 2024) further confirming their phenotype and strongly suggesting that there are multiple subtypes of telocytes in the human DRG (**Fig 1C**).

**Figure 1:**
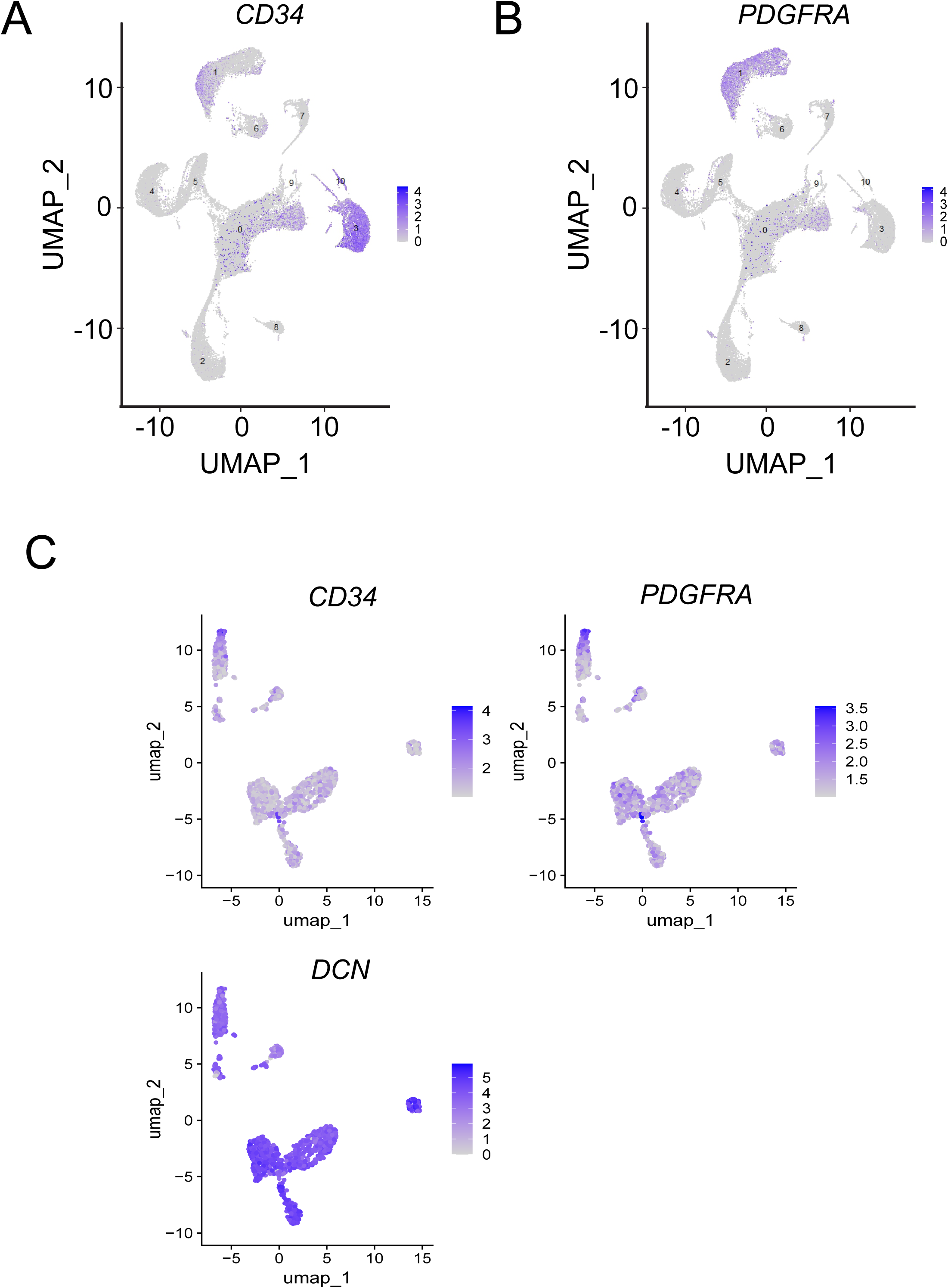
Identification of Telocytes in Human DRG Using Single-Cell Transcriptomics. (A-B) UMAP plots showing the expression of gene markers used to identify telocytes. (C) UMAP plots showing the expression of gene markers that reveal multiple telocyte clusters in the human DRG.

Next, we sought to understand potential interactions between telocytes and human DRG neurons using a ligand-receptors interactome platform (Wangzhou et al., 2021). Looking at ligands potentially produced by telocytes and their interactions with receptors in human DRG we identified 1621 interactions (**Supplementary Table 4**) although many of these were for ligands that have multiple potential receptors. The top 50 ligand-receptor interactions, based on the expression level for the ligand and receptor pair, are shown in **Fig 2A**. The top interaction was the secreted WNT signaling factor *DKK2* from telocytes and the *LRP6* receptor which is expressed in many sensory neuron populations in the human DRG (Bhuiyan et al., 2024; Tavares-Ferreira et al., 2022). Another prominent pair was the *FGF18* gene, which encodes a secreted FGF family protein that can interact with many FGF receptors, in particular *FGFR1* that is highly expressed in human DRG neurons (Bhuiyan et al., 2024; Tavares-Ferreira et al., 2022). Three ligand-receptor pairs indicated a role of telocytes in the interplay between telocytes, neurons and blood vessels. ANGPT1, TNFSF15 and THBS2 represent proteins with strong interaction with neuronal receptors but that have also been shown to maintain vascular structure (Huang et al., 2023; Saharinen & Alitalo, 2011; Zhang & Li, 2012). The telocyte ligand LSAMP belongs to the IgLON family of glycoproteins that are released and after extracellular processing integrated into the matrix of the neuronal cell surface (Kubick, Brosamle, & Mickael, 2018). It has possible interactions with a variety of neuronal receptors.

**Figure 2:**
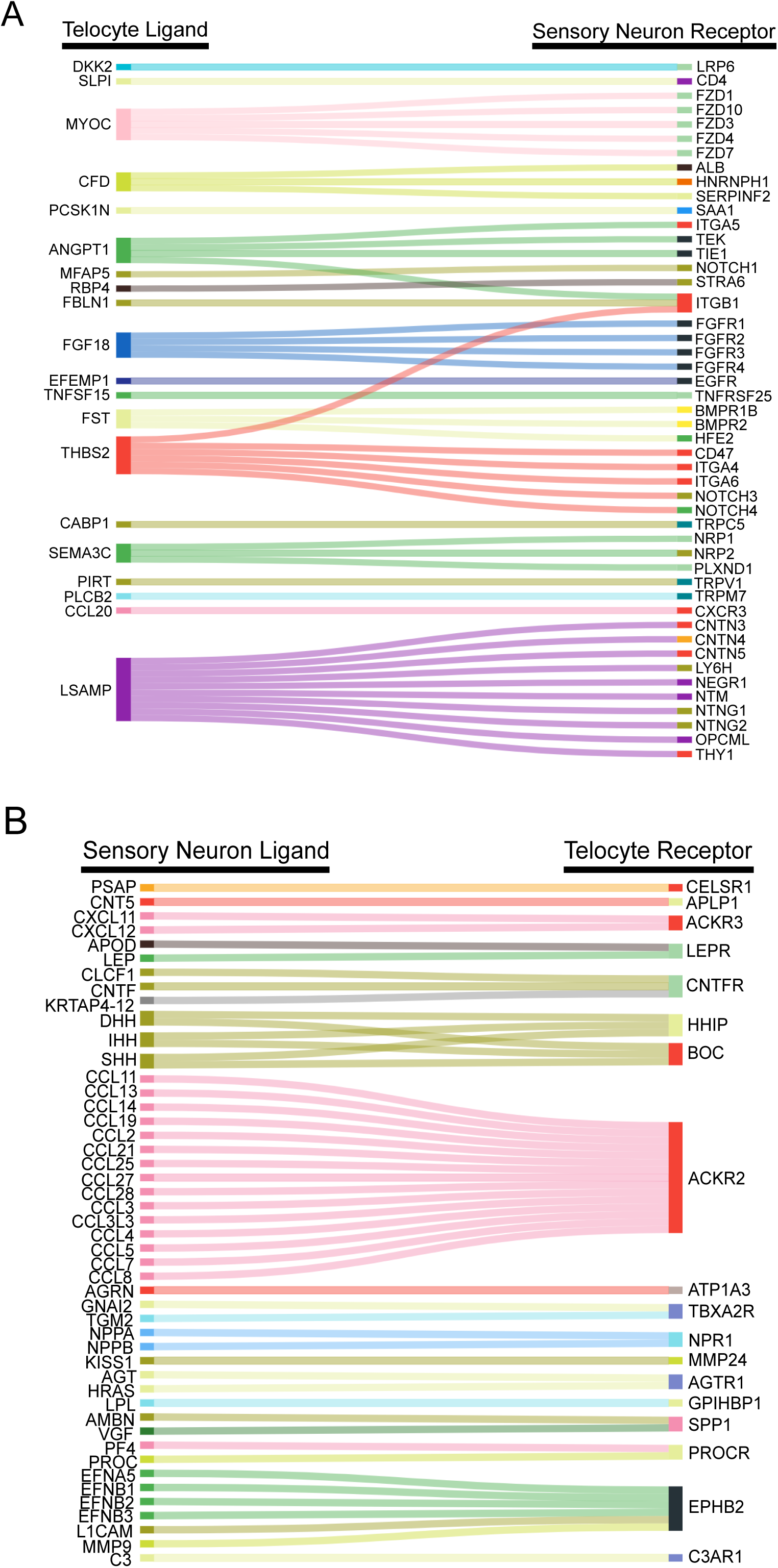
Interactome Analysis Between Neurons and Telocytes in Human DRG. (A) Sankey plot showing the top 50 unique telocyte-to-hDRG ligand-receptor interactions. B) Sankey plot illustrating the top 50 hDRG-to-telocyte ligand-receptor interactions.

The interactome for ligands in human DRG neurons with receptors expressed by telocytes included 1830 interactions (**Supplementary Table 5**) with the top 50 shown in **Fig 2B**. The top ligand-receptor interaction from sensory neurons to telocytes was the *PSAP* gene that encodes prosaponin and the receptor *CELSR1*, which is a member of the cadherin family. We also observed many interactions with chemokines expressed by human DRG neurons and the atypical cytokine receptor type 2 (*ACKR2*) that binds promiscuously to many cytokines to limit their interactions with their cognate receptors. The CNTF receptor complex (*CNTFR*) is an additional cytokine receptor for the CNTF/LIF family of cytokines. CNTF is expressed in neurons and Schwann cells (Hu et al., 2020).

GO enrichment analysis and KEGG-enrichment pathway analysis of the top 150 genes associated with telocyte subclusters showed that characteristic pathways are associated with individual subclusters. One of the nine subcluster showed that 4 out of 6 most enriched molecular functions (with a log10 (observed/expected)) were associated with extracellular matrix and collagen, whereas another telocyte subcluster showed half of the most enriched molecular functions were chemokine related and the vast majority of enriched biological processes was associated with inflammation and immune system activation (**Supplementary Table 6**). A third characteristic subcluster was associated with 21 (out of 32) strongly enriched biological processes that were related to blood vessels and/or vascular endothelium (**Supplementary Table 6**).

### 3.2 Multiple labelling IHC reveals the distribution of CD34+ve cells in human DRG

To gain understanding into the distribution of telocytes in the DRG we used multiple labelling immunohistochemistry which revealed CD34-immunoreactivity in the endothelium of blood vessels, in single mononuclear cells in the interstitial space between neurons, and in cells with the morphology of telocytes with a spindle-shaped cell body and long telopods surrounding the NF200/S100+ve neuron-satellite complex (**Fig. 3**). Telocytes were present close to the S100+ve satellite cell layer surrounding the somata of neurons but they did not extend their telopods between satellite cells or onto the surface of neurons. The number of CD34+ve cells was high around neuron-satellite cell clusters and less prominent in areas with nerve fibre bundles (**Fig. 3**). These CD34+ve telocytes were primarily present in the endoneurium and the space between endo- and perineurium in nerve fibre bundles and in the perivascular space around the microvasculature and around small arterioles (**Fig. 3**). The CD34+ telocytes were negative for CD31. This observation was in contrast with the CD34 immunoreactivity in the vascular endothelium that also showed immunoreactivity for CD31 (data not shown).

**Figure 3:**
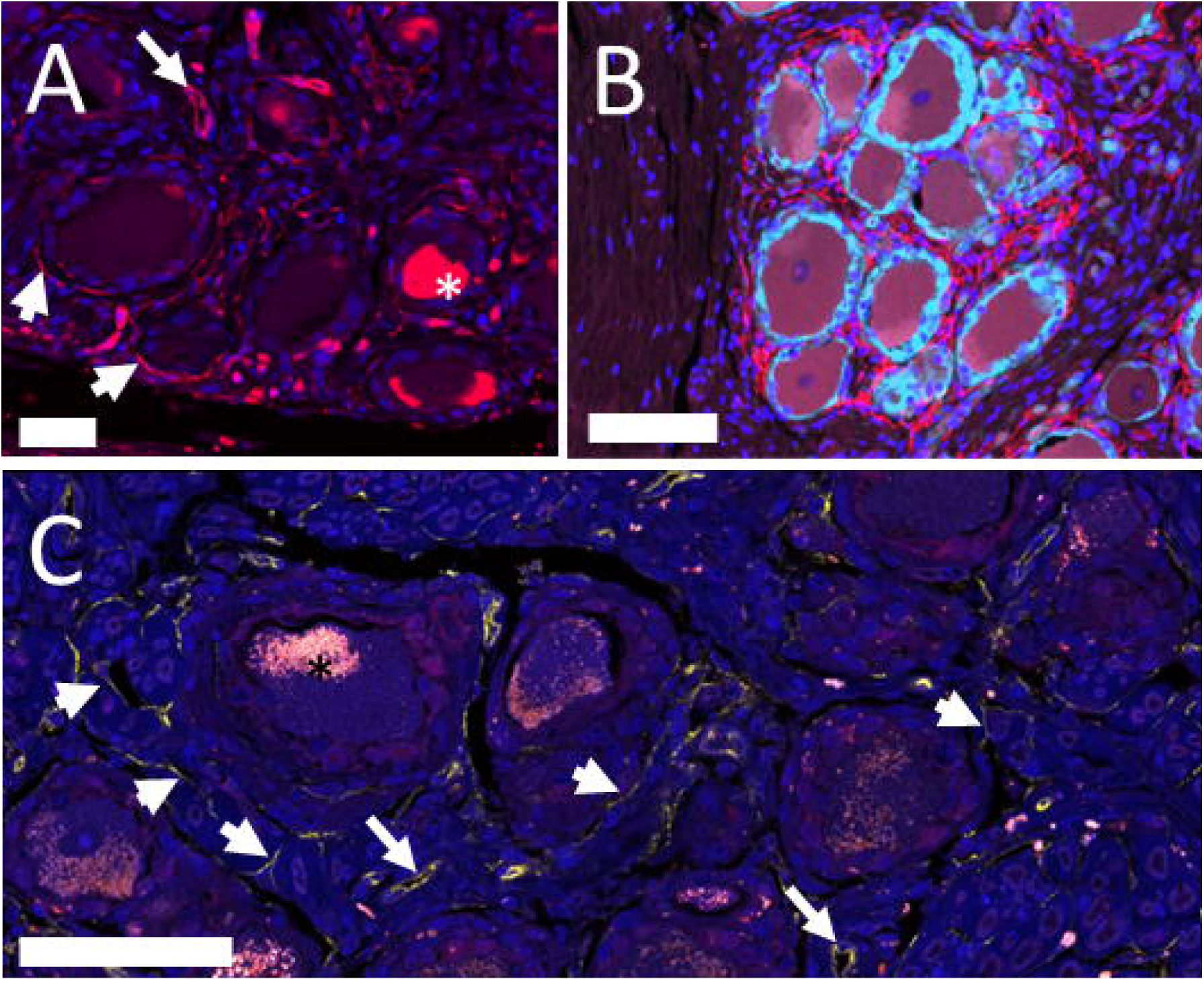
Single and multiple labelling immunohistochemistry human DRG. (A) Immuno-reactivity for CD34 is present in vascular endothelium (long arrow) and in telocytes (short arrows) next to neuronal cell bodies. Nuclei are stained with DAPI. Cell bodies contained autofluorescent lipofuscin (star). Bar = 50 mm. (B) Double labelling with the marker of satellite cells, S100 (pale blue) shows the presence of telocytes and their telopods (red) outside of the neuron-satellite complex. Nuclei are stained with DAPI. Bar = 100 μm. (C) Confocal laser scanning image with a thickness of 1 μm shows the presence of CD34+ fibres (short arrows) next to neuron-satellite complexes, in the endoneural space and next to blood vessels (long arrows). Nuclei are stained with DAPI. Bar = 75 μm.

### 3.3 Transmission electron microscopy outlines ultrastructural features of telocytes in human DRG

Next, to gain more detailed insights into the ultrastructural morphology of the human DRG, we used transmission EM to confirm the presence of telocytes and their networks of telopods. The presence small stellate- or spindle-shaped soma, an oval nucleus with intense heterochromatin and thin, and long telopods were in line with observations of the characteristic telocyte features. The nuclei in DRG telocytes were elongated with dimensions of about 2-4 μm wide and 7-9 μm long and showed heterochromatin at their margins (**Figs. 4, 5**). These telocytes were typically bipolar or tripolar, occasionally featuring areas with slightly enlarged telopod diameters known as podoms, both containing vesicles and caveolae. Telocytes also generally, but not always possessed a basal lamina, often accompanied by extracellular bundles of collagen fibres (**Fig. 5**).

**Figure 4:**
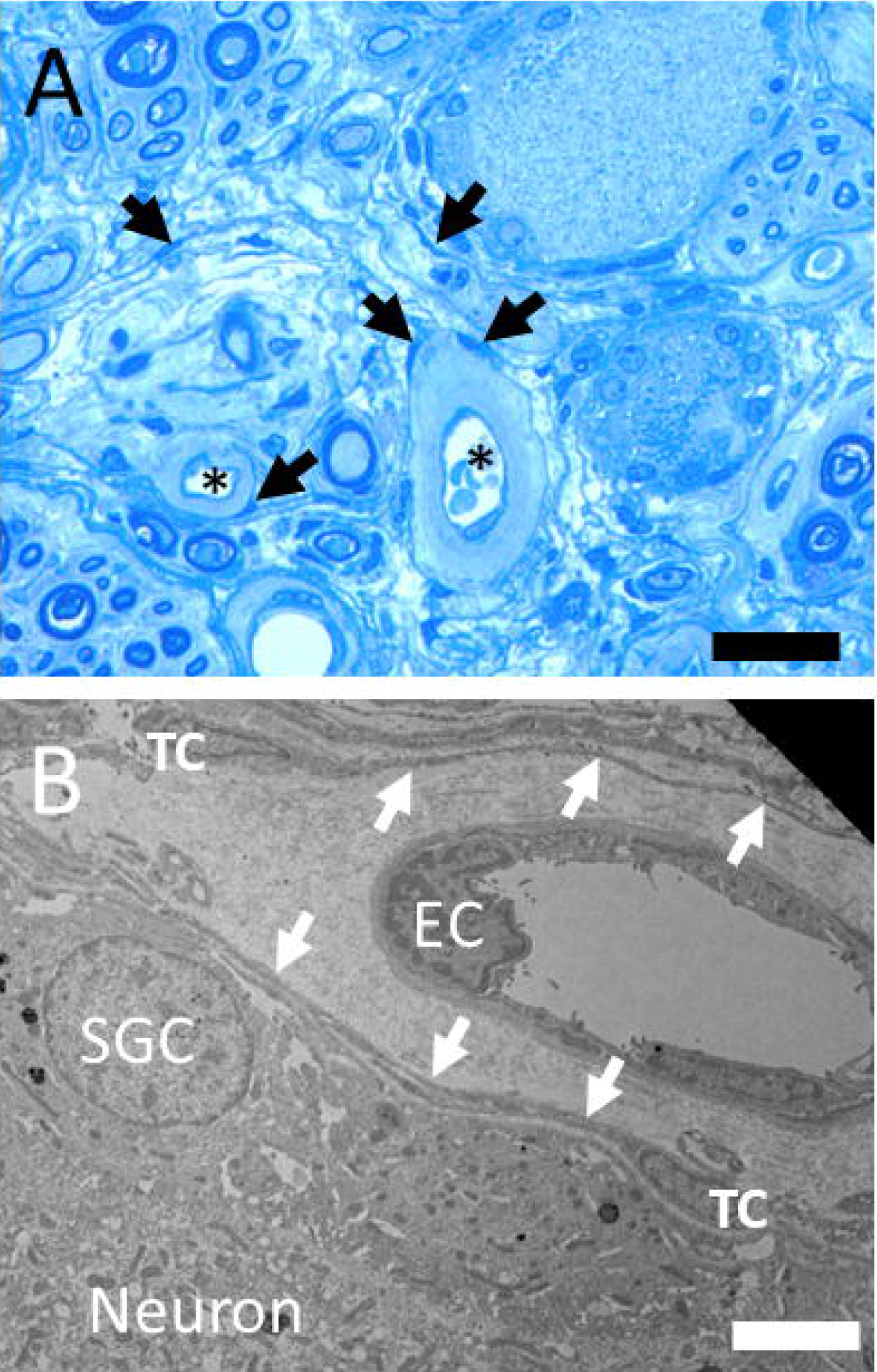
(A) Toluidin blue stained semithin section with neuron-satellie complexes, nerve fibres, capillaries (stars) and pale surrounding perivascular space. Telocyte nuclei in the perivascular and endoneural space are indicated with arrows. Bar = 30 μm. (B) Transmission electron microscope image of a human DRG. Telocyte (TC) with telopods (arrows) between the perivascular space of a capillary (EC – endothelial cell nucleus) and the complex of nerve cell (neuronal soma) and satellite glia cell (SGC). Bar = 4 μm.

**Figure 5:**
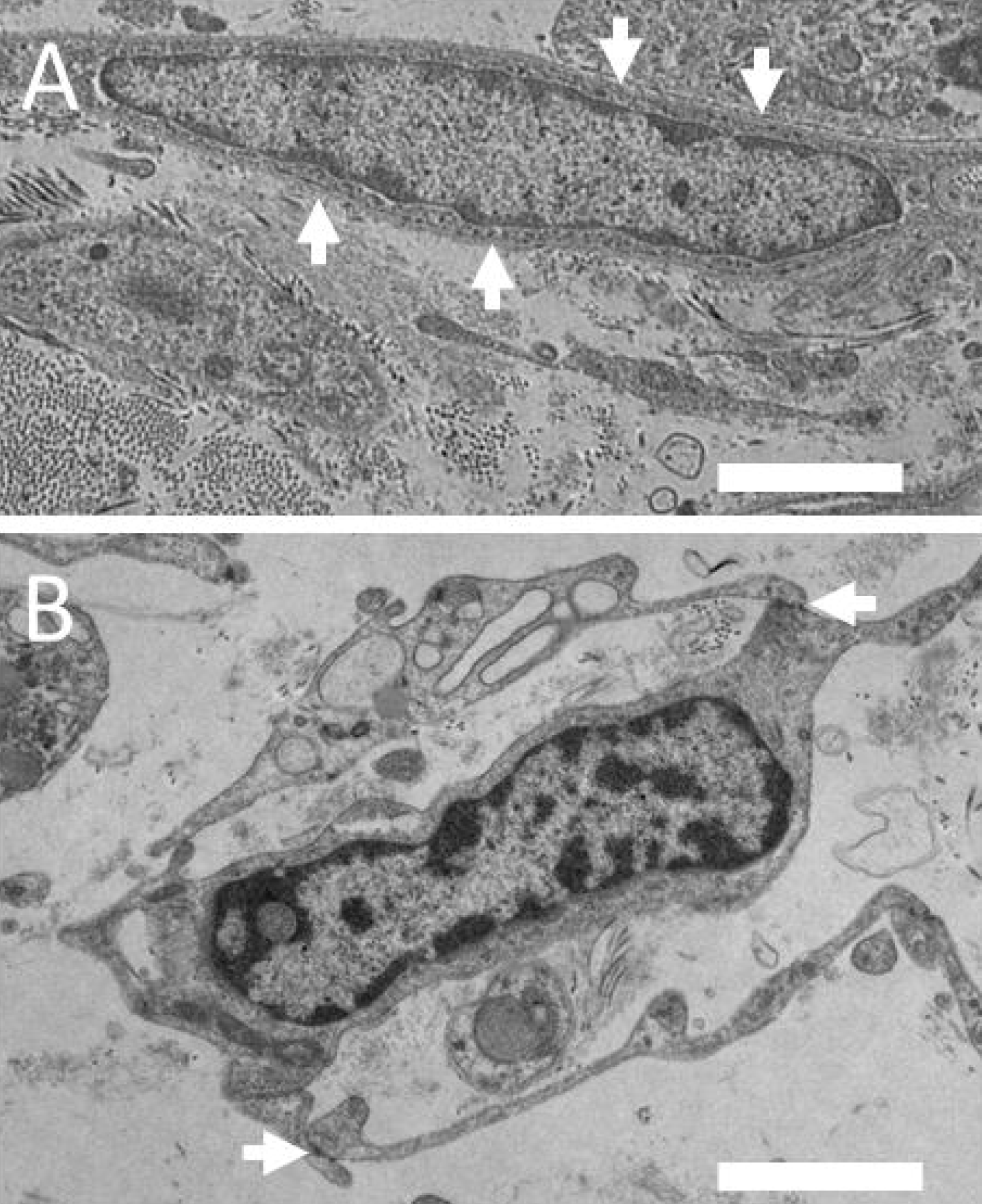
(A) The electron microscopical image shows the elongated nucleus of telocytes, arrow indicate the basal lamina usually associated with soma and telopods of telocytes. Bar = 2 μm. (B) Electron microscopical image of a telocyte with cell-cell contacts indicated with arrows. Bar = 2 μm.

Telocytes were located adjacent to the nerve cell/satellite glia complex (**Fig. 4**) but were also present in the internal perineurium as well as the endoneurium of nerve fibre bundles and in the perivascular space around small blood vessels (**Supplementary Fig. 1**). One DRG specimen showed remnants of exudate around small vessels suggesting the presence of infection of inflammation. Interestingly, the vessel-aasociated telocytes in this sample were bordering the enlarged, exudate-filled perivascular space (**Fig. 4**).

Telopods of perivascular telocytes and telocytes spanning the other parts of the endoneural space were found running parallel to each other without cell-cell contacts. Occasionally electron-dense cell-cell contacts between telopods were observed (**Fig. 5**) but no cell-cell contacts of telocytes with satellite glia cells or neurons could be identified. Telopods of telocytes that surround small blood vessels were usually shorter compared to the extended processes of telocytes that surround nerve fibre bundles or traversed the remaining endoneural space (**Supplementary Fig. 1**).

## 4 Discussion

Most investigations on DRG focus on neurons but the ganglia are comprised of various cell types including satellite cells or endothelial cells which outnumber neurons. However, the identification of cell types in human DRG is primarily based on recent transcriptional studies, which might exclude or underrepresent certain cell types. One cell type that has not been investigated so far are telocytes.

### 4.1 Marker genes and expression patterns define nine subclusters of telocytes

In this study, we used two previously published single nucleus (Bhuiyan et al., 2024) and single cell RNA sequencing datasets (Hou et al., 2024) to identify putative telocytes in the human DRG. Telocytes were identified using the marker genes *CD34* and *PDGFRA*, with *DCN* used as an additional coexpression marker that is less specific for telocytes (Guo et al., 2024). Our analysis revealed that these cells constitute approximately 1.5-3% of all human DRG cells in these datasets.

Reclustering of these cells from the single cell sequencing dataset, which captures cytoplasmic mRNAs giving a more comprehensive view of gene expression in these cells, suggests the presence of several subtypes of telocytes in the DRG. This is supported by studies on the murine intestine where different telocyte subpopulations have been described (reviewed by (Shoshkes-Carmel, 2024)). Our analysis identified nine clusters and subsequent GO enrichment analysis and KEGG-enrichment pathway analysis revealed differential gene and signalling pathway enrichments among these clusters. Each cluster contained specific enrichments in genes and signalling pathways including those associated with immune system, vasculature or connective tissue. Analysis of possible communication between telocytes and neurons revealed numerous potential ligand-receptor interactions between telocytes and sensory neurons, and vice-versa. Notably, the expression of atypical cytokine receptor (*ACKR2*) (Bonavita, Mollica Poeta, Setten, Massara, & Bonecchi, 2016) in telocytes could limit the local action of chemokines released by DRG neurons.

### 4.2 Location and ultrastructure of telocytes

Our analysis of sequencing data indicates the presence of telocytes in human DRG and suggests their potential involvement in various functions. Location of cells is associated with function. Therefore, the location and ultrastructure of telocytes in human DRG was further investigated. Cells expressing CD34 mRNA are known to be present in human DRG and have been reliably identified in other studies using antisera against the CD34 protein (Diaz-Flores et al., 2023; Rosa et al., 2023; Sanches et al., 2023). This study identified CD34 immunoreactive cells in small blood vessels without a muscularis layer, likely capillaries of small venules. These vessels exhibited strong immunoreactivity confined to the endothelium consistent with previous studies in human tissue (Fina et al., 1990; Pusztaszeri, Seelentag, & Bosman, 2006; Rusu, Manoiu, Cretoiu, Cretoiu, & Vrapciu, 2018). Additionally, immunoreactivity for CD34 was present in elongated cells with distinct telocyte-like morphology and long and thin processes. Those cells were located in the endoneural interstitial space adjacent to blood vessels, neuron-satellite complexes and in between nerve fibre bundles. Based on their location, morphology, and CD34 expression, those cells are likely telocytes as corroborated by the description of telocytes in other human tissues (Diaz-Flores et al., 2020; Rusu et al., 2016; Traini et al., 2018).

We then used transmission electron microscopy (TEM) to better understand the characteristic morphology of human DRG telocytes, revealing the ultrastructure of their elongated ovoid nucleus, telopods, podoms and podomers (S. M. Cretoiu & Popescu, 2014; Diaz-Flores et al., 2020; Xiao & Bei, 2016). The size of telocyte cell bodies was consistent with measurements reported in the literature (see review(Wang, Jin, Ma, Zhu, & Wang, 2016)). Telocytes were found in the perivascular space, surrounding microvessels and near neuron-satellite complexes in human DRG similar to their distribution in the human trigeminal ganglion which shares morphological and transcriptional features with the DRG (Bhuiyan et al., 2023; Rusu et al., 2016). The cell bodies and telopods of human DRG telocytes were present in the endoneural space between neuron/satellite complexes and were often, but not always, surrounded by a basal lamina. However, telocytes did not directly connect to the cell bodies or processes of sensory neurons, nor did we find conclusive evidence of any connections with satellite cells. Nonetheless, the proximity of telocytes to neurons, blood vessels and within the endoneural, space supports our expression analysis data which suggests the presence of different telocyte subtypes or functional states.

### 4.3 Connection of expression and location to function

Although this study provides critical insights into the complexity of telocyte biology in the human DRG, however, the precise role of telocytes in human DRG remains unclear. Insights into human telocyte function have been mostly derived from studies using tissue samples from patients. Although functional data on telocytes in humans are limited (Albulescu et al., 2015; Liao et al., 2021; Manole, Cismasiu, Gherghiceanu, & Popescu, 2011; Sanches et al., 2023), descriptive studies showed that the number and localisation of telocytes is altered in different diseases such as atherosclerosis, megaureter or dermal fibrosis (Diaz-Flores et al., 2020; Rosa et al., 2021; Smith et al., 2023; Wishahi et al., 2021; Xu et al., 2021). In this study we identified telocytes throughout the endoneural connective tissue space forming a three-dimensional network in the human DRG. This network may be considered as part of the stroma, providing structural support to the surrounding tissue, given that stromal-cell derived factor-1 has been shown to promote telocyte proliferation (Sanches et al., 2023). However, telocytes might play a more active role in tissue function beyond structural support, especially since the ultrastructure of these cells characterized by abundance of vesicles and occasional multivesicular bodies involved in cell-cell signalling (S. M. Cretoiu & Popescu, 2014; Ratajczak, Ratajczak, & Pedziwiatr, 2016) and clearly supported by the expression analysis findings of this study.

Indeed, telocytes from other tissues are capable of releasing vesicles into the extracellular environment (Fertig, Gherghiceanu, & Popescu, 2014) which interact with neighbouring cells (Cismasiu & Popescu, 2015). For instance, cardiac telocytes have been found to transfer vesicles loaded with microRNAs (miRNA) to local stem cells, and then reacquire the repackaged miRNAs via vesicular endocytosis from the same stem cells (Cismasiu & Popescu, 2015). Given the high density of vesicles and multivesicular cargos within human DRG telocytes, it can be hypothesized that similar bi-directional signalling mechanisms may be ongoing between DRG telocytes and local cells, likely the satellite glia and sensory neurons. Our sequencing data supports that human DRG telocytes are abundant in ligands that can interact with receptors found on satellite glial cells and sensory neurons. One miRNA that has been released from telocytes is miR-21, which infers a protective function on DRG neurons but also contributes to neuropathic pain (Buller et al., 2010; Zhou et al., 2015).

Telocytes and their processes surround endothelial cells and adjacent pericytes of small blood vessels which is corroborated by studies in human heart, gut and parathyroid glands (Diaz-Flores et al., 2022; Liao et al., 2021; Smith et al., 2023). This indicates a role of telocytes related to blood vessel function. It is supported by our analysis of sequencing data which showed a high number of vasculature-related genes in one of the telocyte clusters. Capillaries and venules in DRGs are located near neurons-satellite cell complexes that have a high energy demand. The expression of angiopoietin 1 (*ANGPT1*), thrombospondin 2 and TNFSF15 suggests interactions between telocytes, blood vessels and neurons. Angiopoietin 1 is important for the maintenance of vascular structure and vessel permeability, thrombospondin 2 (*THBS2*) is a matricellular protein that inhibits angiogenesis and tumor necrosis factor superfamily-15 (*TNFSF15*) protein is a cytokine that is known as vascular endothelial growth inhibitor which also inhibits endothelial cell differentiation and proliferation (Bornstein, 2009; Brindle, Saharinen, & Alitalo, 2006; Zhang & Li, 2012).

### 4.4 Summary and outlook

In summary, our analysis of gene expression patterns in telocytes suggest, for the first time, the existence of distinct telocyte subpopulations within human DRG. The unique gene expression profiles not only link to specific cellular functions but also shed light on the multifaceted roles of telocytes in the DRG microenvironment. Supported by multiple-labelling immunohistochemistry and ultrastructural investigations, our findings underscore the potential for critical and diverse functions that human DRG telocytes may possess. This discovery opens new avenues for understanding DRG physiology and potentially offers novel targets for therapeutic interventions in conditions with DRG dysfunction.

## Acknowledgements

We acknowledge Yeunhee Kim, Manager of UTD Genome Center for technical support with library prep for RNA-seq and Pat Vilimas, Christopher Leigh and Jim Manavis, Flinders and The University of Adelaide facilities for their technical support with multiple labelling immunohistochemistry and electron microscopy. The authors thank Anna Cervantes, Geoffrey Funk, and Peter Horton at the Southwest Transplant Alliance for tissue recovery from organ donors. The authors are grateful to the organ donors and their families for their gift.

## Data and materials availability

All data associated with this study are in the paper or the Supplementary Materials. Raw sequencing data are available in the SPARC repository DOI **10.26275/pxwy-sric**

## Ethics approval statement

All human tissue procurement procedures were approved by the Institutional Review Boards at the University of Texas at Dallas. Human lumbar DRGs were procured from three organ donors through a collaboration with the Southwest Transplant Alliance. The use of human tissue was approved by the Office of Research Ethics, Compliance and Integrity, The University of Adelaide.

