## Supplementary Table 6 for "Transcriptomic and histological characterization of telocytes in the human dorsal root ganglion"

Supplemental Table 6: GO enrichment and KEGG pathway analysis

**Immune type** - GO enrichment “Biological Processes”

| GO-term | Description | Count | Strength | FDR | Genes |
| --- | --- | --- | --- | --- | --- |
| GO:2000448 | Positive regulation of macrophage migration inhibitory factor signaling pathway | 2/2 | 2.13 | 0.0118 | CXCR4, CD74 |
| GO:0002540 | Leukotriene production involved in inflammatory response | 2/2 | 2.13 | 0.0118 | ALOX5AP, ALOX5 |
| GO:0071726 | Cellular response to diacyl bacterial lipopeptide | 3/4 | 2 | 0.00072 | CD36, CD14, TLR2 |
| GO:0071727 | Cellular response to triacyl bacterial lipopeptide | 2/3 | 1.95 | 0.0176 | CD14, TLR2 |
| GO:0055096 | Low-density lipoprotein particle mediated signaling | 2/3 | 1.95 | 0.0176 | LPL, CD36 |
| GO:0001660 | Fever generation | 2/3 | 1.95 | 0.0176 | IL1B, IL1A |
| GO:1904149 | Regulation of microglial cell-mediated cytotoxicity | 3/5 | 1.91 | 0.0011 | CX3CR1, TYROBP, SPI1 |

**Immune type** - GO enrichment “Molecular function”

| GO-term | Description | Count | Strength | FDR | Genes |
| --- | --- | --- | --- | --- | --- |
| GO:0004051 | Arachidonate 5-lipoxygenase activity | 2/2 | 2.13 | 0.0491 | ALOX5AP, ALOX5 |
| GO:0005102 | Low-density lipoprotein particle receptor activity | 3/15 | 1.43 | 0.0476 | CD36, STAB1, OLR1 |
| GO:0008009 | Chemokine activity | 8/48 | 1.35 | 7.67x10^-6^ | CCL4, CCL4L2, CXCL2, CXCL8,  CCL14, CCL8, CCL3, CCL3L1 |
| GO:0071813 | Lipoprotein particle binding | 4/28 | 1.28 | 0.0187 | CD36, STAB1, MSR1, LPL |
| GO:0028187 | Pattern recognition receptor activity | 4/28 | 1.28 | 0.0187 | TLR2, CD14, CD36, CLEC7A |
| GO:0019956 | Chemokine binding | 4/29 | 1.27 | 0.0187 | CXCR4, ACKR1, CX3CR1, A2M |
| GO:0042379 | Chemokine receptor binding | 9/71 | 1.23 | 7.67x10^-6^ | CCL4L2, CCL8, CCL4, CCL3, CXCL2, CXCL8, CCL14, CX3CR1, CCL3L1 |

**Immune type** - KEGG pathway analysis

| Pathway | Description | Count | Strength | FDR | Genes |
| --- | --- | --- | --- | --- | --- |
| hsa05144 | Malaria | 12/46 | 1.43 | 3.87x10^-12^ | IL1B, SELE, ITGB2, IL6, CD36, CXCL8, ACKR1, SELP, CSF3, ICAM1, PECAM1, TLR2 |
| hsa04610 | Complement and coagulation cascades | 13/82 | 1.33 | 4.11x10^-11^ | VSIG4, C3AR1, C1QC, C1QA, C1QB, C5AR1, ITGB2, THBD, VWF, A2M, PLAT, PLAUR |
| hsa05134 | Legionellosis | 8/55 | 1.29 | 6.9x10^-7^ | CXCL2, NFKBIA, IL6, TLR2, IL1B, CXCL8, ITGB2, CD14 |
| hsa05143 | African trypanosomiasis | 5/36 | 1.23 | 0.00018 | SELE, ICAM1, IL1B, IL6, PLCB2 |
| hsa05133 | Pertussis | 10/73 | 1.26 | 2.86x10^-8^ | IL6, CXCL8, NLRP3, IL1A, IL1B, ITGB2, C1QA, C1QB, C1QC, CD14 |
| hsa05323 | Rheumatoid arthritis | 11/83 | 1.25 | 9.75x10^-9^ | CCL3L1, CXCL2, IL1A, IL1B, CCL3, IL6, CXCL8, ITGB2, TLR2, ICAM1, FLT1 |
| hsa04061 | Viral protein interaction with cytokine and cytokine receptor | 11/96 | 1.19 | 2.86x10^-8^ | CXCL2, CXCL8, CCL4, CX3CR, CXCR4, CCL4L2, CCL3L1, CCL3, CCL8, IL6, CSF1R |

**Vascular type** - GO enrichment “Biological Processes”

| GO-term | Description | Count | Strength | FDR | Genes |
| --- | --- | --- | --- | --- | --- |
| GO:1903587 | Regulation of blood vessel endothelial cell proliferation involved in sprouting angiogenesis | 4/16 | 1.52 | 0.0027 | GATA2, DLL4, PPP1R16B, MMRN2 |
| GO:0003157 | Endocardium development | 3/12 | 1.52 | 0.0218 | SOX18, SOX17, KDR |
| GO:0043117 | Positive regulation of vascular permeability | 4/17 | 1.49 | 0.0031 | PDE2, BMP6, PtP4A3, TACR1 |
| GO:0001946 | Lymphangiogenesis | 3/13 | 1.48 | 0.0249 | TIE1, SOX18, FLT4 |
| GO:0090051 | Negative regulation of cell migration involved in sprouting angiogenesis | 3/15 | 1.42 | 0.0346 | DLL4, TBXA2R, MMRN2 |
| GO:0060055 | Angiogenesis involved in wound healing | 3/15 | 1.42 | 0.0346 | GATA2, KDR, GPR4 |
| GO:0043114 | Regulation of vascular permeability | 9/48 | 1.39 | 5.6x10^-6^ | PTP4A3, TACR1, CLDN5, PLVAP, CDH5, TEK, PDEA2, BMP6, GPR4 |

**Connective tissue type** - GO enrichment “Biological Processes”

| GO-term | Description | Count | Strength | FDR | Genes |
| --- | --- | --- | --- | --- | --- |
| GO:007223 | Cell proliferation involved in metanephros development | 3/7 | 1.75 | 0.0101 | BMP7, PDGFRB, GPC3 |
| GO:0110011 | Regulation of basement membrane organisation | 3/11 | 1.56 | 0.0225 | LAMA2, NID1, LAMB3 |
| GO:2001046 | Positive regulation of integrin-mediated signaling pathway | 3/13 | 1.48 | 0.0308 | LAMA2, NID1, LAMB1 |
| GO:0070141 | Response to UV-A | 3/14 | 1.45 | 0.0335 | EGFR, MMP2, TIMP1 |
| GO:0032332 | Positive regulation of chrondrocyte differentiation | 4/20 | 1.42 | 0.0068 | SOX5, SOX6, RUNX2, PKDCC |
| GO:2001044 | Regulation of integrin-mediated signaling pathway | 4/21 | 1.4 | 0.0075 | LAMA2, NID1, LAMP1, TIMP1 |
| GO:0061036 | Positive regulation of cartilage development | 5/32 | 1.32 | 0.0026 | SOX5, SOX6, RUNX2, MDK, PKDCC |

**Connective tissue type** - GO enrichment “Molecular Function”

| GO-term | Description | Count | Strength | FDR | Genes |
| --- | --- | --- | --- | --- | --- |
| GO:0030021 | Extracellular matrix structural constituent… | 5/15 | 1.64 | 0.00030 | LUM, DCN, PRELP, BGN, VCAN |
| GO:0048407 | Platelet-derived growth factor binding | 3/11 | 1.56 | 0.0438 | COL3A1, COL1A2, PDGFRB |
| GO:0050840 | Extracellular matrix binding | 8/56 | 1.28 | 6.39x10^-5^ | TGFB1, ECM1, BGN, DCN, NID1, FBLN2, CD248, OLFML2B |
| GO:0005201 | Extracellular matrix structural constituent | 17/131 | 1.23 | 6.39x10^-12^ | LTBP4, LTBP1, FBN2, COL6A2, NID1, LAMB1, LAMA2, FN1, BGN, PRELP, FBLN2, DCN, COL1A2, COL3A1, LUM, TGFB1, VCAN |
| GO:0005518 | Collagen binding | 7/66 | 1.15 | 0.00088 | TGFB1, DCN, NID1, FN1, PCOLCE, LUM, COL6A2 |
| GO:0019838 | Growth factor binding | 10/127 | 1.02 | 0.00013 | LTBP4, LTBP1, COL1A2, EGFR, FGFR2, FGFBP2, PDGFRB, COL3A1, IGFBP7, IGFBP4 |

Connective tissue type - KEGG pathway analysis

| Pathway | Description | Count | Strength | FDR | Genes |
| --- | --- | --- | --- | --- | --- |
| Hsa04512 | ECM-receptor interaction | 7/88 | 1.02 | 0.0025 | COL6A2, COL1A2, LAMB1, FN1, LAMA2, LAMC3, FREM1 |
