## Supplementary figures and images for "Transcriptomic and histological characterization of telocytes in the human dorsal root ganglion"

### Supplementary Figure 1

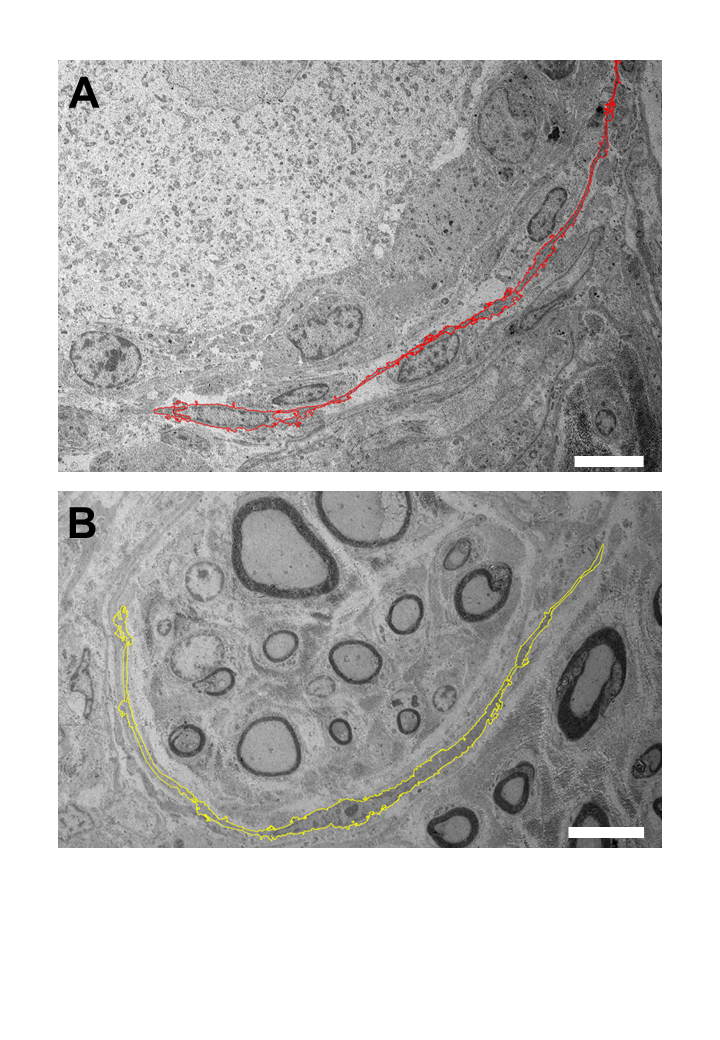
